# Emergence of a recurrent insertion in the N-terminal domain of the SARS-CoV-2 spike glycoprotein

**DOI:** 10.1101/2021.04.17.440288

**Authors:** Marco Gerdol, Klevia Dishnica, Alejandro Giorgetti

## Abstract

Tracking the evolution of the severe acute respiratory syndrome coronavirus 2 (SARS-CoV-2) through genomic surveillance programs is undoubtedly one of the key priorities in the current pandemic situation. Although the genome of SARS-CoV-2 acquires mutations at a slower rate compared with other RNA viruses, evolutionary pressures derived from the widespread circulation of SARS-CoV-2 in the human population have progressively favored the global emergence, though natural selection, of several variants of concern that carry multiple non-synonymous mutations in the spike glycoprotein. These are often placed in key sites within major antibody epitopes and may therefore confer resistance to neutralizing antibodies, leading to partial immune escape, or otherwise compensate infectivity deficits associated with other non-synonymous substitutions. As previously shown by other authors, several emerging variants carry recurrent deletion regions (RDRs) that display a partial overlap with antibody epitopes located in the spike N-terminal domain (NTD). Comparatively, very little attention has been directed towards spike insertion mutations prior to the emergence of the B.1.1.529 (omicron) lineage. This manuscript describes a single recurrent insertion region (RIR1) in the N-terminal domain of SARS-CoV-2 spike protein, characterized by at least 41 independent acquisitions of 1-8 additional codons between Val213 and Leu216 in different viral lineages. Even though RIR1 is unlikely to confer antibody escape, its association with two distinct formerly widespread lineages (A.2.5 and B.1.214.2), with the quickly spreading omicron and with other VOCs and VOIs warrants further investigation concerning its effects on spike structure and viral infectivity.

## 1. Introduction

Coronaviruses generally accumulate mutations at a much lower rate than other RNA viruses, thanks to the efficient proofreading exonuclease activity exerted by nsp14, in complex the activator protein nsp10 (Denison et al., 2011; Ma et al., 2015). As a result, the rate of molecular evolution of SARS-CoV-2 is currently estimated (as of December 5^st^ 2021, based on GISAID data (Shu and McCauley, 2017)), to be close to 25 substitutions/year per genome, i.e. 8.36×10^−4^ substitutions/site/year, which is slightly higher than previous estimates for human endemic coronaviruses (Ren et al., 2015). Consistently with comparative genomics data obtained from other members of the *Sarbecovirus* subgenus, such mutations are not evenly distributed across the genome, but they are disproportionally located in the S gene, which encodes the spike glycoprotein. It is also worth noting that the S gene undergoes frequent recombination events, likely as a result of naturally occurring co-infections in the animal viral reservoirs (Boni et al., 2020), and that these events are also theoretically possible among different SARS-CoV-2 lineages (Ignatieva et al., 2021). The encoded transmembrane protein forms a homotrimer and plays a fundamental role in the interaction between the virus and host cells, promoting viral entry though the interaction with different membrane receptors (Millet et al., 2020). In the case of SARS-CoV-2 and of the closely related SARS-CoV responsible of the 2002-2004 outbreak, such receptor is represented by the angiotensin converting enzyme 2 (ACE2) (Ren et al., 2008; Walls et al., 2020).

While most of these mutations have little or no phenotypic impact at all, some may significantly influence viral transmissibility and the ability of the virus to escape host immune response. The causes underpinning such phenotypic effects may either lie in an increased viral shedding, in the alteration of the binding affinity between the spike receptor binding domain (RBD) and the host ACE2 receptor, or in the modification of key antibody epitopes. The most striking example of a non-synonymous mutation which had a dramatic impact on the dynamics of the pandemics is most certainly represented by S:D614G. This mutation, which was not present in the ancestral lineage that caused the Wuhan outbreak, emerged in the very early phases of the pandemics, quickly becoming dominant worldwide (Korber et al., 2020), most likely due to an increased packing of functional spike protein into the virion (Zhang et al., 2020).

Even though the mutation rate of the SARS-CoV-2 genome remained relatively stable throughout 2020, growing evidence soon started to point out the presence of shared mutations across multiple independent lineages, suggesting ongoing convergent evolution and possible signatures of host adaptation (van Dorp et al., 2020a). While early investigations failed to identify evidence of increased transmissibility associated with such recurrent mutations (van Dorp et al., 2020b), the nearly contemporary independent emergence of three variants sharing the non-synonymous substitution N501Y in the spike protein started to raise serious concerns about the possible involvement of this mutation in increasing viral infectivity. While the functional role of N501Y still remains to be fully elucidated, structural modeling points towards a possible function in the stabilization of the spike protein in the open conformation, which may increase ACE2 binding, especially in combination with other mutations targeting the RBD (Nelson et al., 2021; Teruel et al., 2021; Zhu et al., 2021).

B.1.1.7 (the alpha variant, according to WHO labeling), one of the emerging lineages carrying N501Y, spread in southeastern England in early 2020 and quickly became dominant in Europe. Despite being significantly more transmissible than wild-type genotypes (Davies et al., 2021), alpha was not associated with significant immune escape from the neutralizing activity of convalescent or vaccinated sera (Lustig et al., 2021; G.-L. Wang et al., 2021; Z. Wang et al., 2021; Xie et al., 2021). On the other hand, some point mutations present in the spike NTD, i.e. the deletion of a codon in position 144, led to full escape from the activity of a few NTD-directed monoclonal antibodies (P. Wang et al., 2021).

Two other major lineages carrying N501Y, designated as variants of concerns (VOCs) in early 2021, i.e. B.1.351 (beta) and P.1 (gamma), were linked with major outbreaks in geographical regions with very high estimated seroprevalence, i.e. in the Eastern Cape region (South Africa) (Sykes et al., 2021) and in Manaus (Amazonas, Brazil) (Sabino et al., 2021), respectively. Both variants were characterized by a constellation of non-synonymous mutations and accelerated rates of evolution, which suggested that their selection might have occurred in immunocompromised patients with persistent viral infection (Choi et al., 2020). Among the many features shared by beta and gamma, the most remarkable one was the presence of two additional RBD mutations, i.e. E484K and K417N/K417T. The former one has been identified as a key player in antibody escape, due to its presence in a major epitope recognized by class II RBD-directed antibodies (Greaney et al., 2021; Starr et al., 2020; Starr et al., 2021). On the other hand, mutations of K417, located in an epitope recognized by class I antibodies, are thought to provide a minor contribution to polyclonal antibody response escape (Greaney et al., 2021) and to possibly stabilize, together with E484K and N501Y, the interaction between the RBD and the ACE2 receptor (Nelson et al., 2021). Following these events, the trends of new infections connected with emerging variants started to be subjected to closer monitoring due to the possible negative impact they might have on massive vaccination campaigns (Madhi et al., 2021; Shen et al., 2021).

In the spring of 2021, the lineage B.1.617.2 (delta) was internationally recognized as the fourth VOC. Like the three previously mentioned variants, delta carried several non-synonymous mutations in the S gene, including S:L452R, which is located in a major class III RBD-directed antibody epitope (Shen et al., 2021) and allows to completely escape the neutralizing activity of several monoclonal antibodies (mAbs) (Starr et al., 2021). Following its initial association with the surge of infections that occurred in India in early 2021 (Dhar et al., 2021), this variant rapidly spread worldwide and replaced alpha, which strongly suggested a higher intrinsic transmissibility (Dagpunar, 2021), possibly due to a more efficient cleavage site between the S1 and S2 subunits (Peacock et al., 2021). At the same time, delta was also found to be endowed with significant immune escape properties, which resulted in reduced sensitivity towards the sera of convalescent and vaccinated individuals (Planas et al., 2021) and in reduced vaccine effectiveness, in particular after the first dose (Lopez Bernal et al., 2021). Although delta became dominant worldwide in the second half of 2021, the novel viral lineage B.1.1.529 started to quickly spread in the Gauteng province (South Africa), outcompeting delta. The higher fitness of this lineage has been tentatively linked with a substantial ability to evade immunity from previous infection (Pulliam et al., 2021), which might be consistent with the high number of non-synonymous mutations (32) and indels (three deletions plus one insertion) observed in the S gene compared to the reference SARS-CoV-2 genome. These include a number of previously described RBD mutations associated with the aforementioned VOCs, such as K417N, T478K and N501Y, plus E484A. Based on early epidemiological data, on the growing number of imported cases reported abroad and on the expected immune evasion properties of this variant, on December 1^st^ WHO included B.1.1.529 in the list of VOCs under the “omicron” designation.

Several VOCs and VOIs (including alpha, beta, delta and omicron) carry spike deletions in the NTD. Such deletions were previously shown to often occur in distinct NTD sites, named Recurrent Deletion Regions (RDR), arising in different geographical backgrounds, in independent viral lineages. Some RDR sites display a significant overlap with known immune epitopes, suggesting that they may drive antibody escape (McCarthy et al., 2021). Comparatively, prior to the emergence of omicron, which carries a three amino acids-long insertion (S:ins214EPE) in the NTD of the spike protein, very little attention had been directed towards insertions. Nevertheless, such events are known to have played a fundamental role in the past evolution of SARS-CoV-2 spike protein by allowing, among the other things, the acquisition of a furin-like cleavage site, which is an uncommon feature in bat coronaviruses. This short motif, which is thought to be a key pathogenicity determinant (Johnson et al., 2021), is indeed completely absent in the closely related *Sarbecovirus* RaTG13 (Ge et al., 2016) and only partly present in the recently described RmYN02 (Zhou et al., 2020) and RacCS203 (Wacharapluesadee et al., 2021).

The present work reports the independent occurrence of at least 41 distinct insertion events at the very same NTD site, located between Val213 and Leu216, which will be hereafter referred to as Recurrent Insertion Region 1 (RIR1). The transient international spread of the RIR1 insertion-carrying lineages A.2.5 and B.1.214.2, the presence of S:ins214EPE in the VOC omicron and the identification of several insertions at this site in the delta lineage point out that more attention should be put towards the functional characterization of these codon acquisitions in the near future.

## 2. Materials and methods

### 2.1. Sequence data analysis

The global frequency of insertion and deletion mutations mapped on the SARS-CoV-2 S gene was retrieved, based on GISAID data (Shu and McCauley, 2017), from https://mendel.bii.a-star.edu.sg/ (last accessed on December 5^th^, 2021; credit to Raphael Tze Chuen Lee). Disruptive insertion and deletion mutations (i.e. those that interrupted the open reading frame of the S gene) and insertions carrying undetermined amino acids were discarded. Genomes carrying insertions at any position between codons 213 and 216 were grouped based on the inserted nucleotide sequence. Each group was assigned a code based on progressive Roman numerals, following their chronological order of identification; a lowercase letter was added to design variants of the same insertion that determined a change in the encoded amino acid sequence. The nucleotide sequences of representative entries for each of the identified insertions were aligned with the Wuhan-Hu-1 isolate SARS-CoV-2 reference sequence (GenBank ID: NC_045512.2) using MUSCLE (Edgar, 2004) in the MEGA X environment (Kumar et al., 2018), initially preserving codon boundaries. The multiple sequence alignment was then manually refined to reflect the most probable location of the insertion within each codon and each event was consequently classified as a phase 0, phase I or phase II insertion. All SARS-CoV-2 genome data used for phylogenetic inference in this study were retrieved from GISAID (last access date December 5^th^, 2021) (Shu and McCauley, 2017). In detail, all available sequenced genomes belonging to the lineage A.2.5, to the related sublineages A.2.5.1, A.2.5.2 and A.2.5.3, and to the sister lineage A.2.4 were downloaded, along with associated metadata. While all available GISAID entries were considered for reporting observation frequencies, only high quality genomes (i.e. those listed as “complete” and “high coverage”) associated with a sampling date were taken into account for further analysis. Genomes containing long stretches of Ns (i.e. comprising more than 25 consecutive undetermined nucleotides) were discarded. The reference isolate Wuhan-Hu-1 was also included for tree rooting purposes. Note that several genome sequences from Panama with sampling date anterior to November 2021 were discarded due to the unreliability of associated metadata (i.e. the sampling dates appeared to be inconsistent with the very small genetic distances with recent isolates belonging to the same lineage). Overall, the A.2.5-focused datasets included 1,283 sequences.

SARS-CoV-2 genomes were analyzed with the nextstrain *augur* pipeline (https://github.com/nextstrain/augur). Briefly, nucleotide sequences were aligned with MAFFT (Katoh et al., 2002) and the resulting multiple sequence alignment was used as an input for a maximum likelihood phylogenetic inference analysis, carried out with FastTree (Price et al., 2010) under a generalized time reversible model of molecular evolution. The resulting tree was further refined in the *augur* environment with treetime v.0.8.1 (Sagulenko et al., 2018) using sampling date metadata, generating a time-calibrated tree. The phylogenetic tree were rooted based on the oldest available genotype, which in this case was Wuhan-Hu-1, and graphically rendered using FigTree v.1.1.4.

A root-to-tip genetic distance analysis was performed by plotting the sampling dates against the total number of nucleotide substitutions (excluding insertions and deletions) observed in genomes belonging to the A.2.5 lineage and related sublineages. These were calculated with MEGA X (Kumar et al., 2018), compared with the reference genotype Wuhan-Hu-1. The global average genome-wide mutation rate of SARS-CoV-2, roughly equivalent to 25 substitutions per year, was retrieved from GISAID (as of December 5^th^, 2021).

### 2.2. System setup of coarse-grained models

The simulations on the wild-type spike protein were carried out considering the crystallographic structure deposited in PDB (accession ID: 6XR8) (Cai et al., 2020). A few missing portions were modeled with Swiss Model (Waterhouse et al., 2018) in order not to compromise the molecular dynamic properties of the protein. Homology modeling was performed to obtain the 3D structure of the spike protein of A.2.5 with Swiss Model (Waterhouse et al., 2018), using the EPI_ISL_1502836 GISAID entry as a reference. The protein structure was converted to a coarse-grained Martini representation using the martinize.py script (Aguayo-Ortiz et al., 2017). The coarse-grained protein coordinates were then positioned in the center of a simulation box of size 23×23×23 nm^3^.

The Martini coarse-grained force field with an Elastic Network (CG-ElNeDyn) (Aguayo-Ortiz et al., 2017) was used for running the molecular dynamics simulations through the Gromacs 2019.3 package (Van Der Spoel et al., 2005). The analyses were run using isothermal-isobaric NPT ensemble equilibrium simulations. The temperature for each group (protein, water and ions) was kept constant at 315K using V-rescale thermostat (Bussi et al., 2007) with a coupling constant of 1.0 ps. The pressure was isotropically controlled by a Parrinello-Rahman barostat (Martonák et al., 2003) at a reference of 1 bar with a coupling constant of 12.0 ps and compressibility of 3×10^−4^. Non-bonded interactions were used in their shifted form with electrostatic interactions shifted to zero in the range of 0-1.1 nm. A time step of 20 fs was used with neighbor lists updated every 20 steps. Periodic boundary conditions were used in the x, y and z axes. ∼4μs were collected for the simulations of the wild type and mutant (i.e. A.2.5) spike proteins, respectively. The root mean square deviation of backbone beads (RMSD), the root mean square fluctuations (RMSF) and the radius of gyration (RGYR) were calculated using the gmx rms, rmsf and gyrate modules from the Gromacs package (Van Der Spoel et al., 2005). Principal component analysis (PCA), computed with MDAnalysis, was restricted to backbone beads, as it is less perturbed by statistical noise and provides significant characterization of the essential space motions (Michaud-Agrawal et al., 2011). To visualize the direction and extent of the principal motions of the simulated systems, a porcupine plot analysis was performed using the modevectors.py script in Pymol (DeLano, 2002).

## 3. Results and Discussion

### 3.1. Presence of a recurrent insertion region (RIR1) in the N-terminal domain of SARS-CoV-2 spike protein

The analysis of the genomic data deposited in GISAID as of December 5^th^, 2021 revealed that S gene insertions (excluding those that disrupted the open reading frame) were present in just a minor fraction of all sequenced SARS-CoV-2 genomes, i.e. roughly 0.3% of the total. Prior to the emergence of omicron, the impact of insertions in the S gene on viral evolution had been very limited compared with deletions, which are found in several widespread VOCs and VOIs, such as alpha, beta and delta. Overall, the frequency of observation of spike deletions was more than 500 folds higher than spike insertions, even though this ratio is expected to significantly change in the future, along with the spread of omicron. As previously reported by other authors, most deletions occur in specific sites of the N-terminal domain, including the four previously identified Recurrent Deletion Regions (RDR) 1, 2, 3 and 4 (**Figure 1**) (McCarthy et al., 2021) and the deletion which characterizes the delta variant, occurring at positions 157/165. This is consistent with the higher rate of mutation observed for the S1 region (which includes the NTD and RBD) in human coronaviruses compared with the more slowly evolving S2 subunit (Kistler and Bedford, 2021). Despite their lower frequency of observation, insertions do not occur randomly in the S gene. In fact, the overwhelming majority of the insertion mutations mapped so far in SARS-CoV-2 S gene target the NTD, being in most cases (nearly 3,000 genomes) identified at a specific site, located between codons 213 and 216 (**Figure 1**). However, this figure might be an underestimate due to the frequent use of reference-based insertion-unaware algorithms for SARS-CoV-2 genome assembly, especially during the early phases of the pandemics. Due to the convergent finding of such insertions in independent viral lineages (see below), this region will be hereafter named Recurrent Insertion Region 1 (RIR1). Even though insertions were observed at several other spike sites, RIR1 was the only one where multiple insertions have independently occurred in different lineages. The only spike insertions site displaying a higher absolute number of sequenced SARS-CoV-2 genomes (as of December 5^th^, 2021) is S:ins145T, found in the VOI mu (Laiton-Donato et al., 2021) (**Figure 1**).

**Figure 1.**
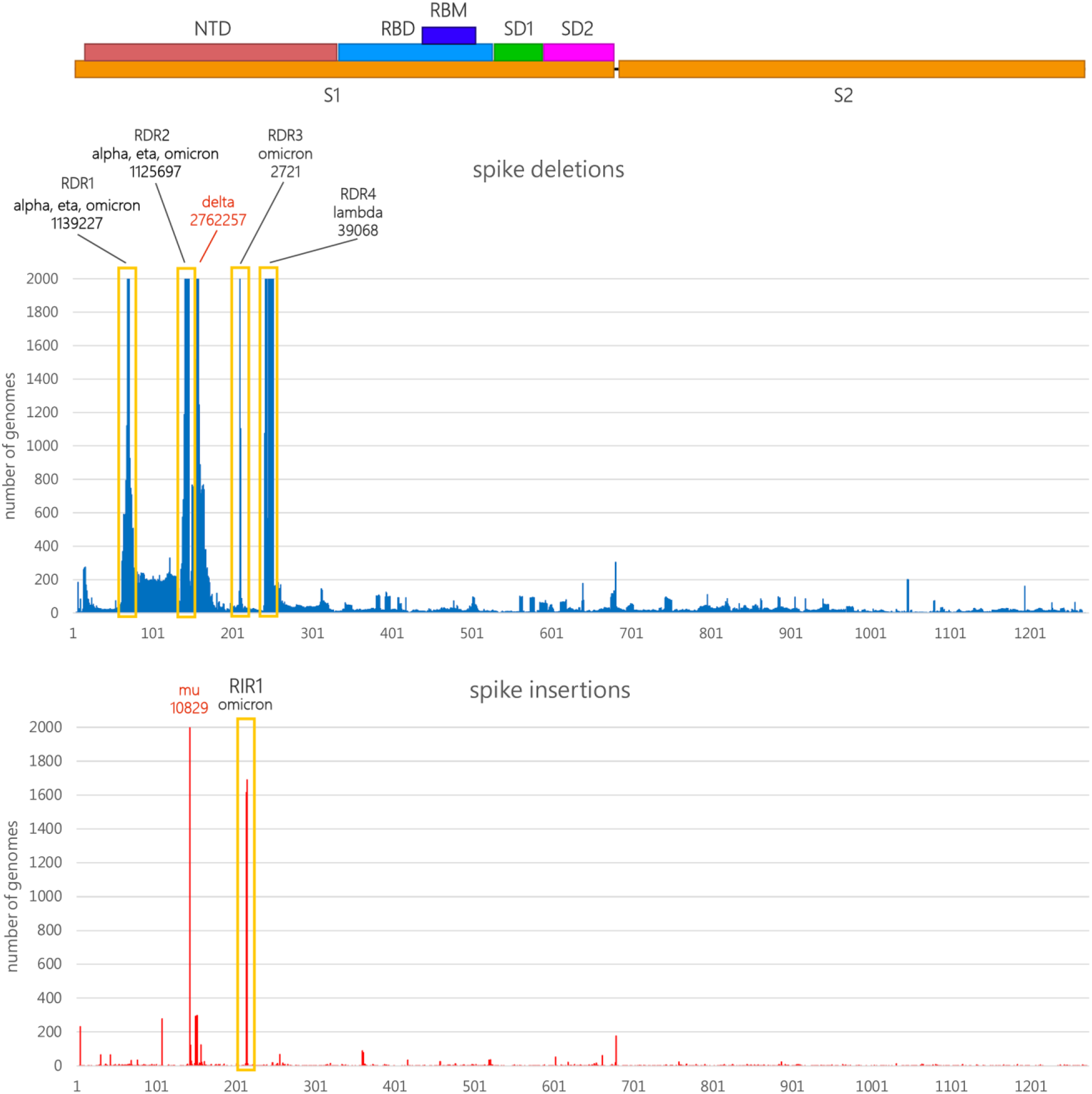
Schematic representation of the SARS-CoV-2 protein, with indication of the two functional S1 and S2 subunits, which are separated by a furin-like proteolytic cleavage site, the N-terminal domain (NTD), the receptor binding domain (RBD) and receptor binding motif (RBM), the SD1 and SD2 subdomains. The absolute number of observed deletion mutations along the S-gene are reported (https://mendel.bii.a-star.edu.sg/ was last accessed on December 5^th^, 2021). Bars were truncated at 2,000 observed genomes; in such cases, the absolute number of observations is reported above the truncated bars, together with the main VOCs and VOIs associated with each indel. The position of RDR1-RDR4 from a previous study (McCarthy et al., 2021), as well as the deletion 157/158 characterizing the delta variant and the ins145T insertion characterizing the mu variant, are reported.

### 3.2. RIR1 insertions independently emerged in multiple viral lineages

As of December 5^th^, 2021 RIR1 insertions could be documented as the result of at least 41 independent events that occurred in different branches of the SARS-CoV-2 phylogenetic tree, which strongly suggests convergent evolution. Even though the length of the insertion spanned from one to eight codons (**Figure 2**), the overwhelming majority of the genomes with RIR1 insertions (99% of the total) only included three codons (**Table 1**). While most RIR1 insertions were associated with very small local clusters that not lead to further spread, a few were found in viral lineages with widespread community transmission. These include the lineages B.1.1.529/omicron (insertion XXXIX), A.2.5 (insertion III), B.1.214.2 (insertion IV) and also B.1.639 (insertion IX), which was never associated with a high number of infection but displayed continuous circulation for several months (**Table I**). While omicron is currently the subject of intense scrutiny by the scientific community, A.2.5 and B.1.214.2 were never included in the list of VOIs, despite reaching high prevalence in some geographical regions. While the former lineage will be discussed in detail as a case study in the following sections, it is worth briefly mentioning here the transient spread of B.1.214.2 in early 2021. Indeed, this lineage was likely imported from central Africa to Europe in late 2020. Here, between March and April 2021, it accounted for a non-negligible fraction of the covid-19 cases recorded in Belgium and Switzerland. Following an initial spread, which led to over 1,000 documented infections worldwide, the frequency of observation of B.1.214.2 started to drop in parallel with the rise of alpha, and further declined when delta became dominant. This lineage has not been detected since early July 2021 and can be thus provisionally considered as extinct. Insertion IV, which results in the addition of the TDR tripeptide between R214 and D215, was associated with the presence of two other spike non-synonymous mutations located on the RBD (i.e. Q414K and N450K). These have been previously linked with a moderately increase in RBD stability (Teng et al., 2021) and immune escape both towards a few mAbs and towards convalescent sera (Liu et al., 2021), respectively. Due to the lack of functional data, it is presently unknown whether insertion IV and the other aforementioned mutations endowed this lineage with improved transmissibility or with increased potential for reinfection.

**Figure 2.**
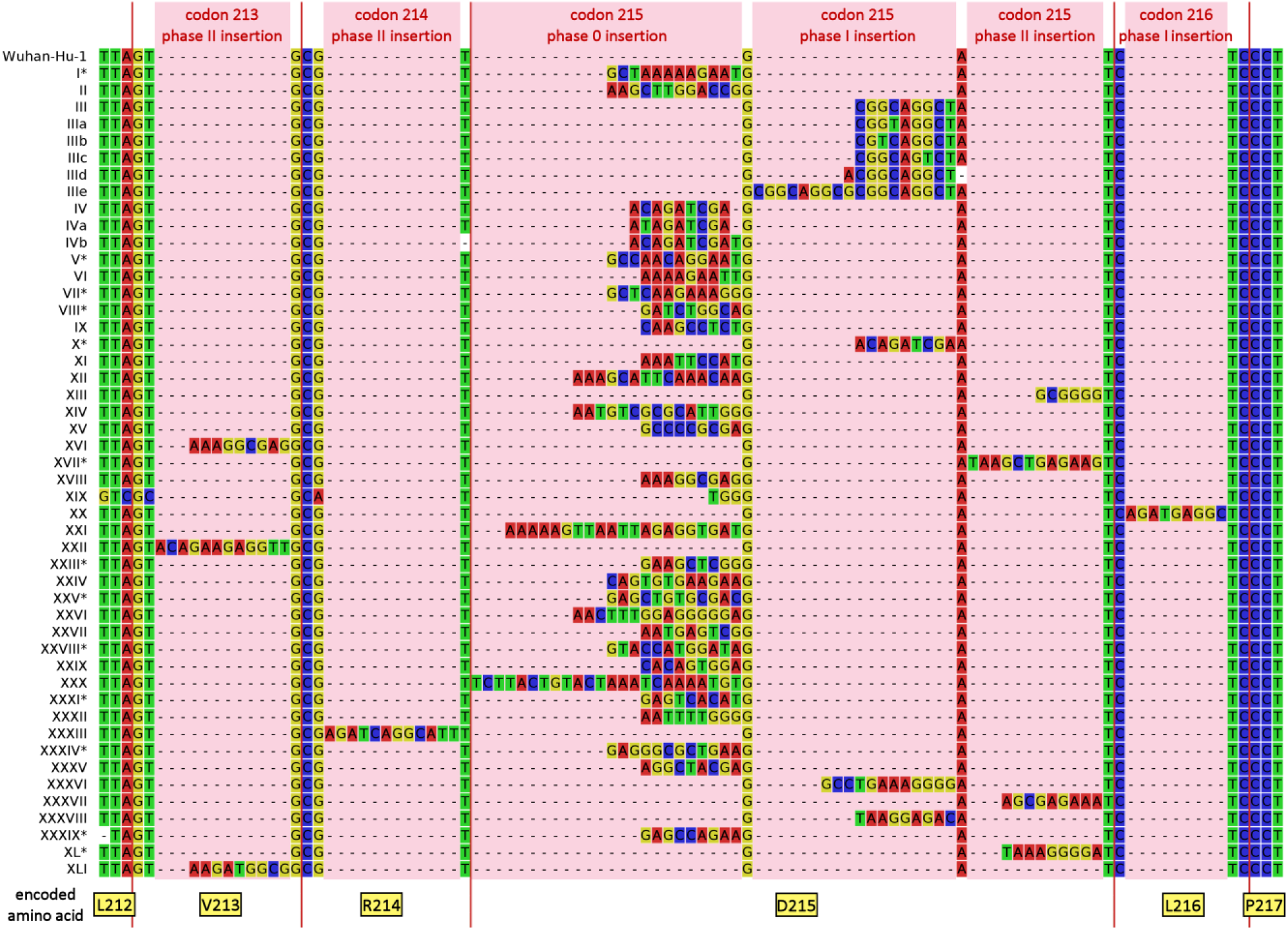
Multiple sequence alignment of the nucleotide sequences of the SARS-CoV-2 S gene of the viral lineages characterized by an insertion at RIR1, compared with the reference sequence Wuhan Hu-1. The multiple sequence alignment only displays a small portion of the S gene and of the encoded spike protein, zoomed-in and centered on RIR1. Red vertical bars indicate codon boundaries, with the encoded amino acids (in the Wuhan Hu-1 reference sequence) indicated below. See **Table 1** for the amino acid sequences encoded by the insertions. Please note that the exact position of all insertion could not be unambiguously detected in all cases; those with ambiguous placement are marked with an asterisk.

**Table 1.**
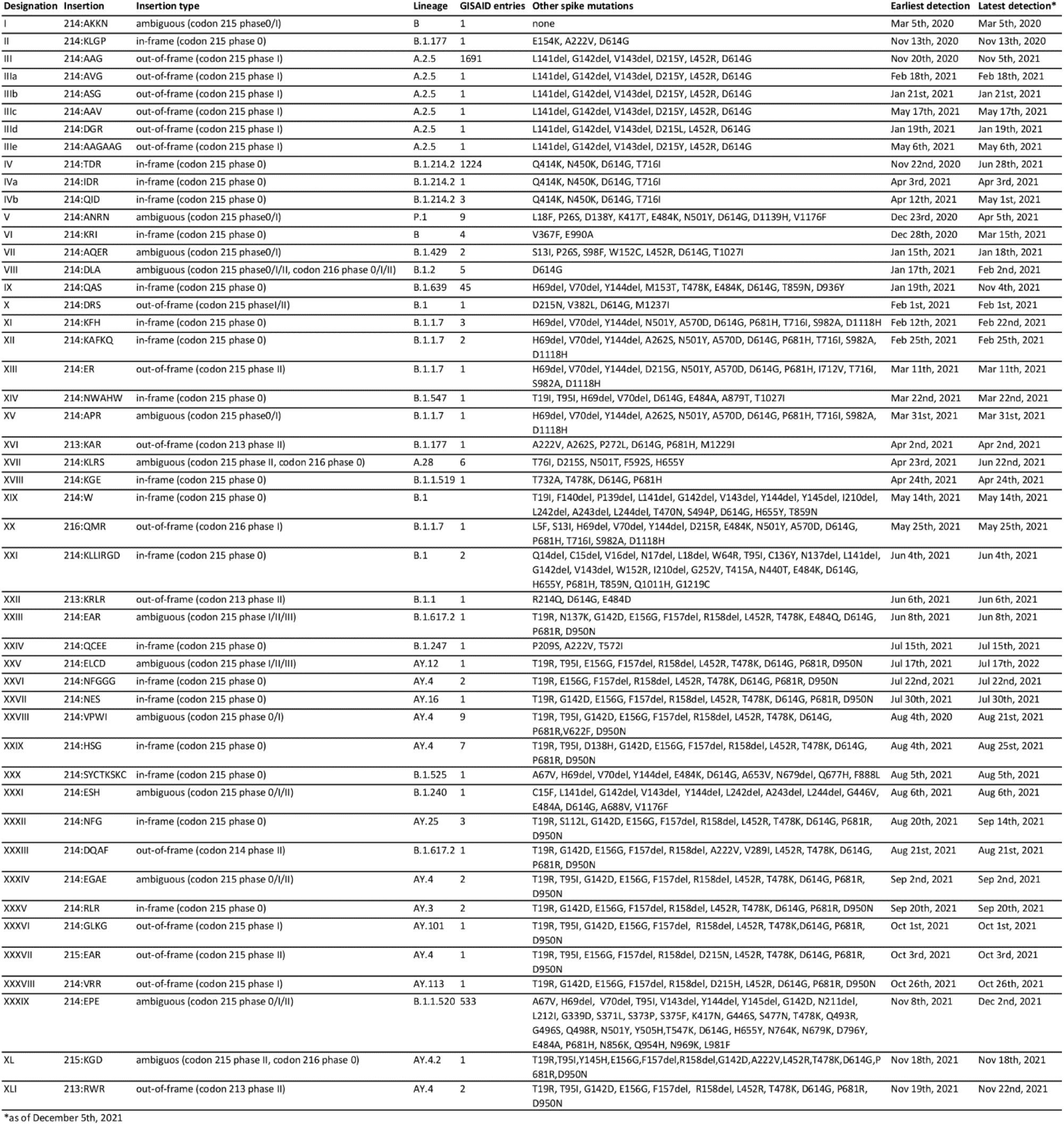
Summary of the 41 independent RIR1 insertions found in the SARS-CoV-2 genome, ordered by the earliest date of detection, as of December 5^th^, 2021.

Several other insertion events at RIR1 occurred in lineages identified as VOCs or VOIs by WHO, CDC, ECDC or PHE, including some that have been recently de-escalated to the status of variants under monitoring. In detail, insertion V was found in eight viral genomes belonging to the gamma lineage, sequenced in different Brazilian states and Guyana between December 2020 and April 2021, indicating the presence of community transmission in the region. As reported in a previous work (Resende et al., 2021), these genomes belong to a monophyletic P.1-like clade that appears to be basal to P.1. The highly transmissible alpha lineage, which became dominant in Europe and quickly spread worldwide in early 2021 (Volz et al., 2021), before the rise of delta, was associated with at least five independent insertions at RIR1 (insertion XI, XII, XIII, XV and XX) between February and May 2021 (**Table 1, Figure 2**). A single RIR1 insertion (XXX) was recorded in August 2021 in the lineage B.1.525 (eta) in the United Kingdom (Ozer et al., 2021) and two sequence genomes characterized by the presence of insertion VII belonging to B.1.429 (epsilon) (Zhang et al., 2021) were sequenced in California in January 2021. Several recently identified insertions at RIR1 are associated with delta (i.e. insertion XXIII, XXV, XXVI, XXVII, XXXII, XXXIII, XXXIV, XXXV, XXXVI, XXVII, XXXVIII, XL and XLI), but none of these have led to significant community spread to date, even though a couple of them were linked to small clusters in England (**Table 1, Figure 2**).

One of the most recent RIR1 insertions (insertion XXXIX) was associated with the emergence of the B.1.1.529/omicron lineage, first detected on November 8^th^, 2021, but possibly spreading undetected in Southern Africa since early October. While the number of omicron cases confirmed by genome sequencing as of December 5^th^, 2021 is still relatively low (see **Table 1**), the rapid spread of this variant worldwide strongly suggest that S:ins215EPE will become the most frequently observed RIR1 insertion in the matter of a few days.

Albeit not directly linked with variants designated as VOCs or VOIs, other RIR1 insertions were associated with the presence of immunologically relevant spike mutations. This is the case of the previously mentioned insertion IX (lineage B.1.639), which is characterized by the contemporary presence of E484K, T478K and by the deletions Δ 69/Δ 70 (found in RDR1) and Δ 144 (found in RDR2), which are shared by several VOCs and VOIs. Curiously, like the omicron insertion XXXIX, insertions XIV, XIX, XXI, XXII and XXXI also targeted viral genomes carrying non-synonymous spike mutations at E484 and deletions at RDR1, suggesting a possible role of RIR1 insertions in compensating otherwise slightly deleterious mutations, like previously hypothesized for RDR1 (Meng et al., 2021).

Taking into account the limited efforts carried out by several countries in genomic surveillance throughout 2020 and 2021, the insertions reported in **Table 1** and **Figure 2** may just represent a fraction of those that emerged at RIR1 during the course of the pandemics. Although it was possible to unambiguously ascertain the exact placement of just 27 out of the 41 RIR1 insertions (see **Table 1**), most of them (17 out of 27 cases, i.e. 63%) were in-frame, occurring at phase 0 between codons 214 and 215, with no effect on neighboring codons. However, others were out-of-frame, occurring either at phase I (i.e. between the first and the second nucleotide of a codon) or at phase II (i.e. between the second and the third nucleotide of codon) (**Figure 2**). In detail, three insertions were observed at phase II within codon 213, one at phase II within codon 214, three and two at phase I and II, respectively, within codon 215, and a single one at phase I within codon 216. In such cases, the placement of the insertion often determined a non-synonymous mutation of the residues flanking RIR1 either at the N- or at the C-terminal side (**Table 1**). It is also worth noting that insertion XXXIX (B.1.1.529/omicron) was associated with a three nucleotides-long, out-of-frame proximal deletion, which affects codons 211 and 212, resulting in the deletion of a single amino acid. Similar deletions were uncommon in other lineages carrying RIR1 insertions, as they have been previously observed in a single other case, i.e. insertion XXI, which displays a Δ 210 deletion.

Although the origins of the 41 RIR1 insertions was not investigated in the present study, other authors have previously suggested that they may be result either from the incorporation of distal regions of the SARS-CoV-2 genome itself, of host mRNAs (Peacock et al., 2021), or even of the genomic sequence of endemic coronaviruses co-infecting the host (Venkatakrishnan et al., 2021). These events would be most likely explained by a still poorly understood copy-choice recombination processes occurring during viral genome replication (Chrisman et al., 2021; Garushyants et al., 2021). Nevertheless, we caution that the short length of RIR1 insertions (usually nine to twelve nucleotides) is in most cases not sufficient to unequivocally establish the origins of the inserted nucleotide sequence, since several randomly occurring identical sequence matches are expected to be found in a broad range of living organisms. On the other hand, when RIR1 insertions are relatively long, the robustness of such inferences might be significantly higher (Peacock et al., 2021).

### 3.3. Mutational pattern of A.2.5 lineage

As mentioned above, the only two lineages carrying insertions at RIR1 with solid evidence of widespread community transmission are A.2.5 and B.1.214.2. The inserted amino acid sequence found in A.2.5 is AAG, as the result of the phase I out-of-frame insertion of the nucleotide sequence CGTCAGGCTA within codon 215, which determines the non-synonymous substitution of Asp215 to Tyr (as a result of a GAT->TAT codon replacement). In a few cases the inserted sequence was duplicated (IIIe), contained a non-synonymous mutation (IIIa, IIIb and IIIc), or was subject to more complex rearrangements (IIId) (**Table 1, Figure 2**). Besides the insertion at RIR1, A.2.5 also displays the deletion of three codons (Δ 141-143) in RDR2 (**Table 1**), sometimes extending to codon 144. This region has been previously implicated in antibody escape (McCarthy et al., 2021) and shows deletions in some relevant VOCs and VOIs, including alpha and eta. In particular, Δ 144 appears to largely explain the resistance towards several NTD-directed mAbs displayed by alpha *in vitro* (Wang et al., 2021). Moreover, the insertion at RIR1 is also combined with L452R, a key mutation that confers resistance towards class III RDB-directed antibodies (Greaney et al., 2021), including LY-CoV555, the basis for the formulation of the commercial mAb bamlanivimab developed by Eli Lilly (Starr et al., 2021). Among the lineages currently or previously designated as VOCs and VOIs, L452R is also found in delta, kappa and epsilon. Like the overwhelming majority of the variants circulating in 2021, A.2.5 is characterized by the presence of the prevalent mutation D614G. Although no other spike mutations are widespread in A.2.5, the A.2.5.3 sublineage acquired S477N, also found in omicron and known to strengthen the binding with the ACE2 receptor (Singh et al., 2021). Overall, this mutation is associated with ∼2% of all A.2.5 genomes (**Figure 3A**). Other relevant spike non-synonymous mutations, known to significantly alter either ACE2 binding or antibody recognition, were only seldom detected: K417T, N501Y and E484K (which are the hallmark spike mutations of gamma) were simultaneously found in a single genome (EPI_ISL_2305075, see **Figure 3A**) sequenced in Texas in May 2021. E484K was found in three additional cases (two in the United States, one in Canada) in April 2021, and N501Y was detected in nine additional cases (eight in the United States, one in Canada) between March and May 2021. Interestingly, in one such cases N501Y was paired with E484Q, which is found in kappa and determines reduced antibody sensitivity, even though not synergistically with L452R (Ferreira et al., 2021). The acquisition of the mutation P681H, known to increase the efficiency of the furin-like cleavage site was documented in 8 cases, 6 of which also displayed N501Y. Such insertions occurred independently in different branches of the A.2.5 evolutionary tree, indicating convergent evolution (see **Figure 3A**). Other lineage-defining non-synonymous mutations of A.2.5 are placed in other genomic locations. These included K1657E, F3071Y, T3255I (shared with delta and mu) and H3580Q in ORF1a; P1000L in ORF1b; S74F and G196V in ORF3a; S197L and M234I (shared with iota) in N (see **Figure 3B**). The functional consequences of these point mutations are presently unknown.

**Figure 3.**
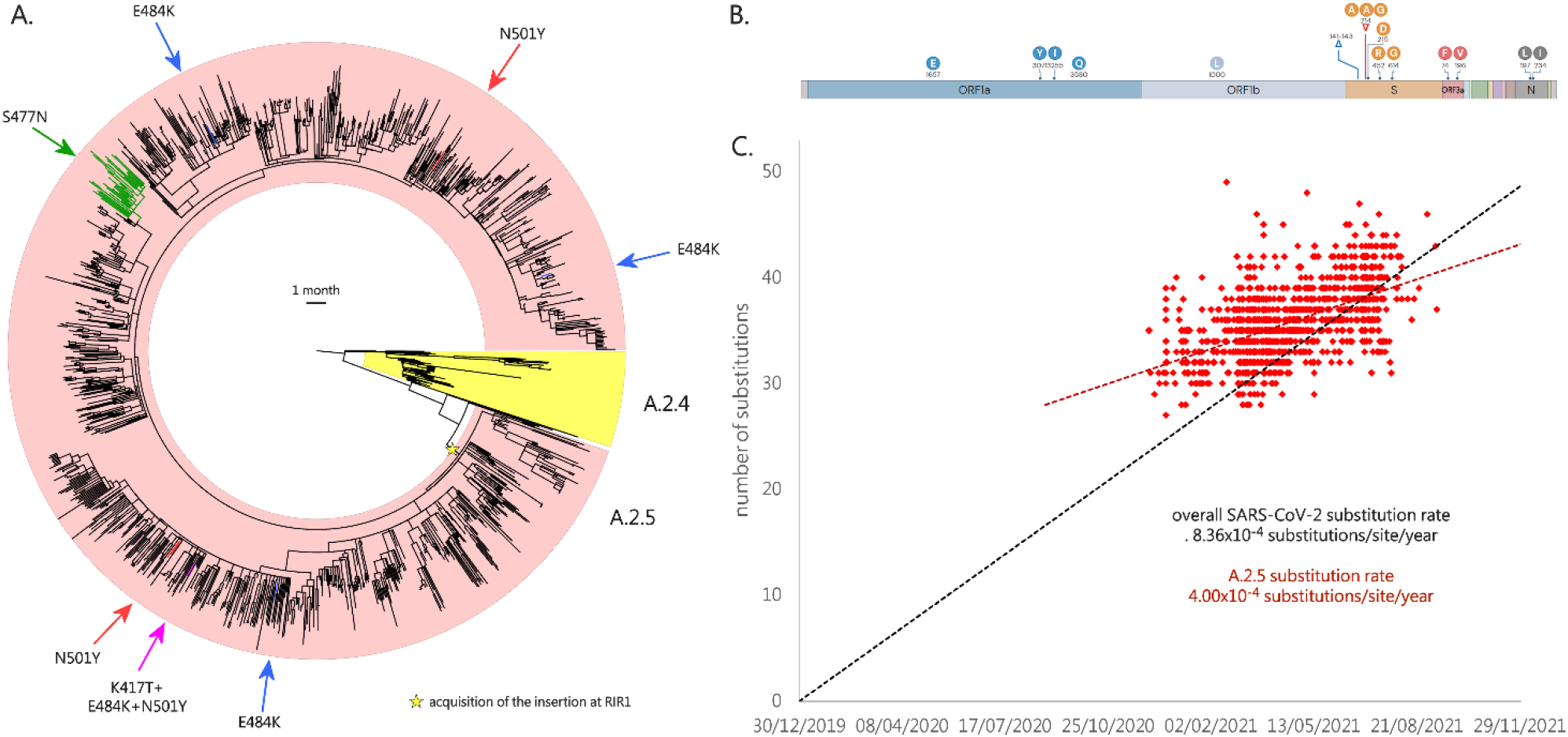
Panel A: circular time tree exemplifying the phylogeny of the A.2.5 lineage related sublineages. Only high quality, complete genomes have been included. The Wuhan-Hu-1 strain was used to root the tree; the sister lineage A.2.4 is also indicated. The acquisition of relevant spike mutations placed in the receptor binding domain (i.e. S477N, K417T, E484K and N501Y) is marked with arrows. Please note that the monophyletic clade linked with the acquisition of S477N corresponds to the A.2.5.3 sublineage. Panel B: key mutations associated with the A.2. lineages. Genes associated with mutations (compared with the reference strain Wuhan-Hu-1) are indicated; only mutations detected in > 50% of the genomes belonging to this lineage and associated sublineages are shown. Modified from https://outbreak.info/. Panel C: root-to-tip genetic distance (number of nucleotide substitutions) of the genomes belonging to the A.2.5 lineage and related sublineages, compared with the reference genome Wuhan-Hu-1. The black dashed line represents the average rate of mutation of all SARS-CoV-2 sequenced genomes, according to GISAID (i.e. 25 substitutions per genome per year, as of December 5^th^, 2021). The red dashed line represent the rate of mutation computed for A.2.5. Note that insertions and deletions were excluded from this calculation.

Root-to-tip genetic distance analysis revealed that the overall nucleotide substitution rate observed in the A.2.5 lineage (and related sublineages) was significantly lower than the average substitution rate computed for SARS-CoV-2 (based on GISAID data, last retrieved on December 5^th^, 2021), as evidenced by the markedly different slope of the regression line (see **Figure 3C**). This was consistent with a substitution rate equal to 4.00×10^−4^ substitutions/site/year, i.e. roughly 12 substitutions/genome/year. Nevertheless, the A.2.5 SARS-CoV-2 genomes detected in the earliest phases of the spread of this lineage (i.e. December 2020) were linked with a number of substitutions significantly higher than the average number of substitutions found in the same period in other SARS-CoV-2 lineages (i.e. ∼33 vs ∼25).

### 3.4. Emergence and international spread of A.2.5

A.2.5 belongs to one of the very few surviving children lineages of the ancestral lineage A, which, after several months of limited global spread, has led to a few major clusters of infections in 2021, such as the one which involved A.23.1 in Uganda (Bugembe et al., 2021). A.2.5 stems from A.2.4, the dominant lineage in the Panama pandemics during the first half of 2020 (Franco et al., 2020). The first documented cases can be traced back to late November 2020, all within a 100 km^2^ area around the capital city Panamá. However, the precise timing of the emergence of A.2.5, along with the acquisition of insertion III at RIR1 and of the other associated mutations described in the previous section, is presently unclear due to the insufficient molecular surveillance carried out in Central America. To date, less than 1,300 out of nearly 480K covid-19 cases reported in Panama have been selected for viral characterization by sequencing i.e. less than 0.3% of the total, far below of the threshold that would be sufficient to track emerging variants (Vavrek et al., 2021). The presence of a number of genomes sampled in El Salvador and Guatemala, two countries where genomic surveillance has been virtually non-existing in 2020, in the earliest-branching clade belonging to A.2.5 (**Figure 3A**), leaves the precise geographical origins of this lineage unclear.

Nevertheless, A.2.5 undoubtedly underwent expansion in Panama between December 2020 and February 2021, as revealed by the increase in estimated prevalence from ∼60 to ∼95%. Interestingly, A.2.5 has been linked with clinically documented reinfections in individuals previously infected by the A.2.4 lineage, which is consistent with the presence of the constellation of non-synonymous spike mutations reported in the previous section, some of which may have immune escape properties (Díaz et al., 2021). The A.2.5 lineage likely spread very early also in the neighboring countries: while investigations carried out in August 2020 failed to identify A.2.5 in Costa Rica (Molina-Mora et al., 2021), the prevalence of this lineage in the country reached 30% between March and June 2021, with the establishment of large clusters of community transmission (**Figure 4**). A.2.5 may have undergone a similar spread in other countries in central America, including Belize, Honduras, El Salvador, Guatemala and Mexico, where multiple cases have been detected, starting from the spring of 2021 (**Figure 4**).

**Figure 4.**
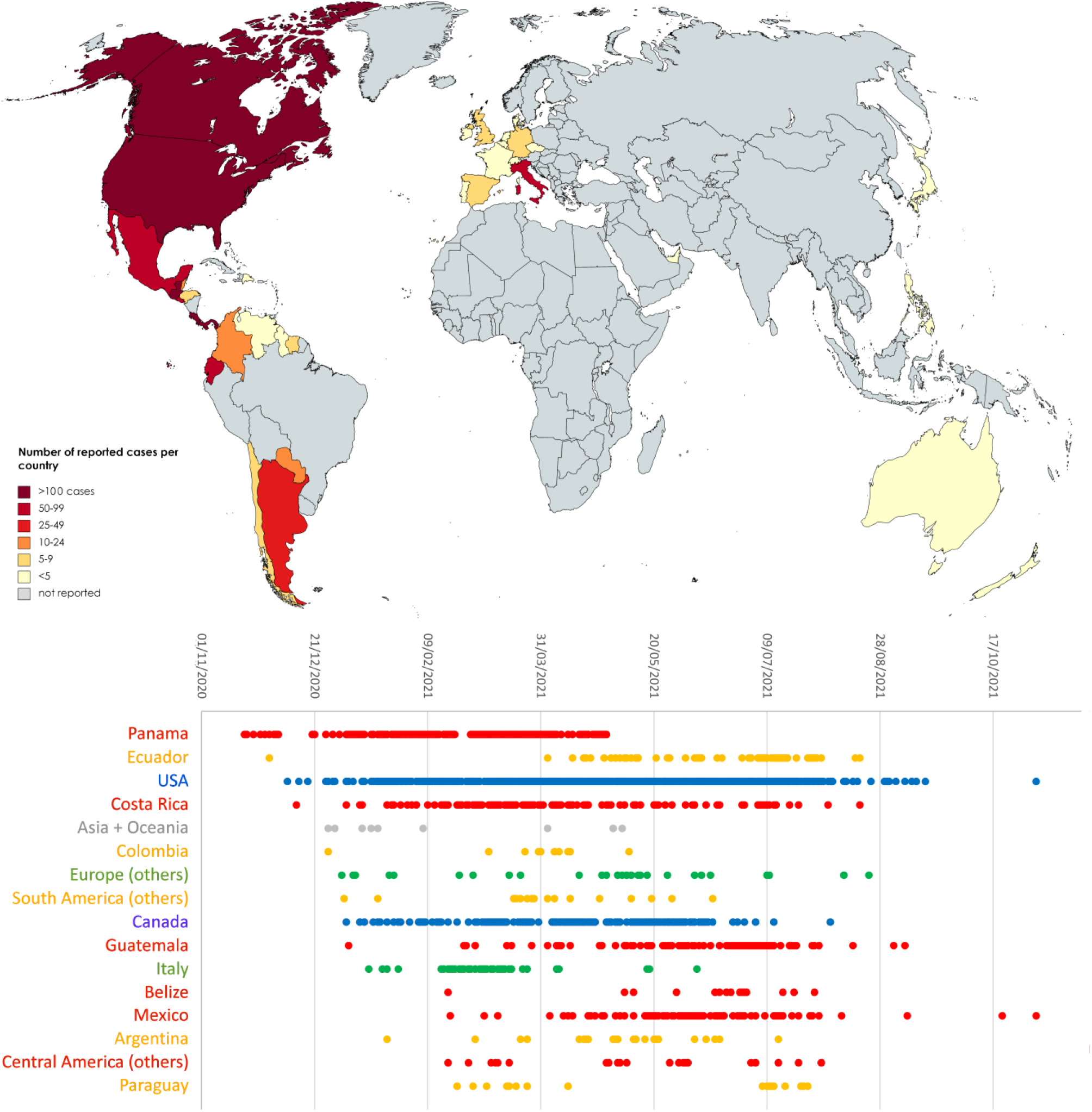
Upper panel: global spread of A.2.5 and related sublineages. Lower panel: detailed timing of the detection of sequenced genomes belonging to A.2.5 and related sublineages in different countries. Only countries with >= 10 unique days of detection are reported, whereas the others were collapsed in geographic macroareas (i.e. Asia + Oceania, Europe, South America and Central America). The reported dates refer to the dates of sampling reported in GISAID.

The remarkable spread of SARS-CoV-2 in Central America was connected with a significant number of exported cases, which have sometimes led to clusters of infection abroad. The first evidence of the detection of A.2.5 in southern America dates to December 1^st^ 2020, in Ecuador. In this country, the acquisition of the spike mutation S477N, mentioned in the previous section, later led to the establishment of the A.2.5.3 sublineage (**Figures 3 and 4**). Reports in other Latin American countries remain sporadic, but it is worth noting that A.2.5 genomes have been so far sequenced in Argentina, Suriname, Guyana, Grenada, Dominican Republic, Sint Marteen, Cayman Islands, Chile, Colombia, Venezuela and Paraguay (**Figure 4**). The earliest cases exported in other continents were reported with similar timing in UAE (December 27^th^, 2020), Philippines (December 30^th^, 2020) and Australia (January 11^th^, 2021), which is consistent with the period with the highest incidence of covid-19 infections documented in Panama. Cases linked with A.2.5 in Europe were identified in Luxembourg, Portugal, Germany, Italy United Kingdom, Czech Republic, France, Belgium, Ireland, Switzerland, Denmark, Netherlands and Spain. In most cases these did not lead to significant community transmission, with the exception of the cluster of cases linked with the A.2.5.2 sublineage recorded in Campania (central Italy) in February-March 2021 (**Figure 4**). Similarly, imported cases have most certainly led to local cluster of infections in different areas of the United States and Canada, starting from late 2020 (**Figure 4**). Nevertheless, the prevalence of A.2.5 in Northern America never exceeded 0.5%). No SARS-CoV-2 infections linked with A.2.5 have been identified to date in the African continent.

The global frequency of observation of A.2.5 and related sublineages underwent a rapid decline in the second half of 2021, in parallel with the global spread of delta. As of December 5^th^, 2021, just five genomes belonging to this lineage have been sequenced over the past three months (**Figure 4**). Further molecular surveillance are expected to reveal whether A.2.5 will disappear, in a similar fashion to what occurred for B.1.214.2 during the summer of 2021.

### 3.5. Impact of RIR1 insertions on the structure of the spike glycoprotein

RIR1 is located in a loop which connects the spike NTD β strands 15 and 16, a region which, unlike RDR1, 2, 3 and 4, does not show any overlap with any known major NTD antigenic sites (Cerutti et al., 2021; McCallum et al., 2021). Hence, the involvement of the insertions reported in this manuscript in antibody escape is unlikely, even though the possibility that this modification may lead to paired structural alterations at distantly related sites, leading to a reduced surface accessibility of canonical antibody epitopes cannot be ruled out. Moreover, the possibility that RIR1 insertions might significantly affect T-cell epitopes, considering that the majority of T-cell response appears to be directed towards the spike NTD and CTD domains (Tarke et al., 2021), cannot be ruled out. Comparative genomics investigations carried out on other viruses belonging to the *Sarbecovirus* subgenus revealed that the RIR1 has been previously prone to structural alterations during the radiation of bat coronaviruses (Garry et al., 2021). In fact, the spike proteins RmYN02, RacCS203, BANAL-116 and BANAL-247 (Temmam et al., 2021; Zhou et al., 2021, 2020), which are among the closest known relatives to SARS-CoV-2 when genomic recombination is taken into account (Lytras et al., 2021), comprise an insertion of four codons at RIR1 in comparison with other bat coronaviruses.

Most certainly, the spread of the A.2.5 and B.1.214.2 lineages in different geographical contexts between late 2020 and early 2021, as well as the recent rapid global emergence of omicron, suggest that RIR1 insertions are unlikely to have a detrimental impact on the three-dimensional structure of the spike protein or to significantly reduce the infectivity of these variants. At the same time, the well-defined length of the insertions (in the overwhelming majority of cases 3 or 4 codons) suggests that some critical structural constraints, that may prevent the selection of shorter insertions or limit their associated evolutionary benefits, might exist. Several spike mutations located in the NTD can affect the structural organization of the spike protein, altering the stability of the interaction between the RBD and the ACE2 receptor, or its accessibility to antibody recognition. For instance, the NTD Δ 69/Δ 70 deletion, which, like RIR1, is found in multiple independent lineages, does not determine a significant antibody escape *in vitro* (P. Wang et al., 2021). However, it is thought to have an important impact on the structure of the spike protein, by conferring increased cleavage at the S1/S2 site, thereby allowing higher incorporation in the virion (Meng et al., 2021). In light of these observations, some NTD indels apparently not related with immune escape may act as permissive mutations, by compensating small infectivity deficits associated with other RBD mutations (i.e. L452R in A.2.5, Q414K and N450K in B.1.214.2, K417N, N440K, G446S, S477N, T478K, E484A, Q493R, Q498R, N501Y and Y505H in omicron).

Interestingly, the insertion of a seven amino-acid long peptide at RIR1 in SARS-CoV-2 through passages in Vero cell cultures has been recently implicated in enhanced *in vitro* infectivity, which may be linked with an increase in the positive charge of NTD surface. According to the authors, this insertion (which is not reported in Table 1 due to its laboratory origin) might have increased the affinity of the spike NTD to heparin, bringing viral particles in close proximity with host cells, thereby favoring the interaction with ACE2 (Shiliaev et al., 2021). While RIR1 insertions rarely share significant pairwise similarity both at the nucleotide and at the amino acid level (**Figure 2, Table 1**), we tested whether the amino acids found in the 41 RIR1 insertions were over-represented compared to expectations (assuming that no codon usage bias was present). As shown in **Supplementary Figure 1**, the basic amino acids arginine and lysine were the most abundant ones (each one accounting for over 10% of total observations), followed by alanine, glutamic acid and glycine. Overall, lysine was the amino acid characterized by the highest observed/expected ratio (i.e. > 3) and also arginine showed a moderate increase in frequency compared to expectations, supporting the conclusions by Shiliaev and colleagues about the benefits of acquiring basic residues at RIR1 for viral infectivity. Nevertheless, the negatively charged glutamic acid was the second most highly enriched amino acid and aspartic acid was also characterized by a positive observed/expected ratio, which may extend such benefits to all charged residues. On the other hand, several amino acids with hydrophobic side chains (e.g. I, M, T, V and Y in particular) were strongly under-represented. In general, however, none only one out of the three widespread lineages with insertions at RIR1, i.e. B.1.214.2, included a positively charged amino acid in the inserted tripeptide, suggesting that the hypothesis proposed by Siliaev and colleagues might not be applicable to all insertions occurring at RIR1. To preliminarily investigate the impact of RIR1 insertions on the structure of the spike protein, we applied molecular dynamics simulations, a well-known technique able to capture and study the dynamical properties of proteins and to assess the effects of mutations, deletions and insertions (Hansson et al., 2002). In this case, we have used a coarse-grained force-field to compare the structural and/or dynamical differences between the spike proteins from the wild-type virus and from the A.2.5 lineage. After 2μs microseconds of simulations, the RMSD of the backbone atoms relative to the equivalent initial structures (which represents a global measure of protein fluctuations) was calculated as a function of time to evaluate the stability of MD simulations equilibrium in the two systems. No significant global displacement was detected for any of the three protein models compared with the initial structure, as most of the RMSD values only displayed fluctuations in a range between 0.35Å and 0.45Å. Similarly, the presence of a few spike mutations in A.2.5 only led to minor changes in the compactness of the protein, as suggested by the differences of about 1 Å found in the average RYGR values among the two models (**Supplementary Figure 2**). On the other hand, some fluctuations were visible in the RMSF of the A.2.5 spike protein model, in particular in the regions which harbored non-synonymous mutations compared with the wild-type protein. To understand changes in the direction of motions of the two systems under analysis, PCA was performed on the last 2μs of the simulations, the time after which the systems reached the equilibration state. The analysis was then restricted to the backbone beads, as they are less perturbed by statistical noise, providing at the same time a significant characterization of the essential space motions (Michaud-Agrawal et al., 2011). The diagonalization of the covariance matrix of fluctuations of the residues belonging to the backbone resulted in a set of eigenvalues, which were plotted in decreasing order against the corresponding eigenvector indices. The first few eigenvectors corresponded to concerted motions that quickly decreased in amplitude to reach some constrained and more localized fluctuations. Here we present the principal modes along the first eigenvector (**Figure 5A**), which covers about 25% of the motions of the protein. Consistently with the placement of non-synonymous mutations (**Figure 3B**), this analysis revealed that A.2.5 exhibited some changes in the fluctuations in regions belonging to the NTD and RBD, which are shown in red in **Figure 5B**. This indicates that the presence of the mutations and insertions may induce local structural and dynamical changes on the spike protein, highlighting the usefulness of performing studies on the dynamical properties of insertions upon their emergence in variants with widespread circulation.

**Figure 5:**
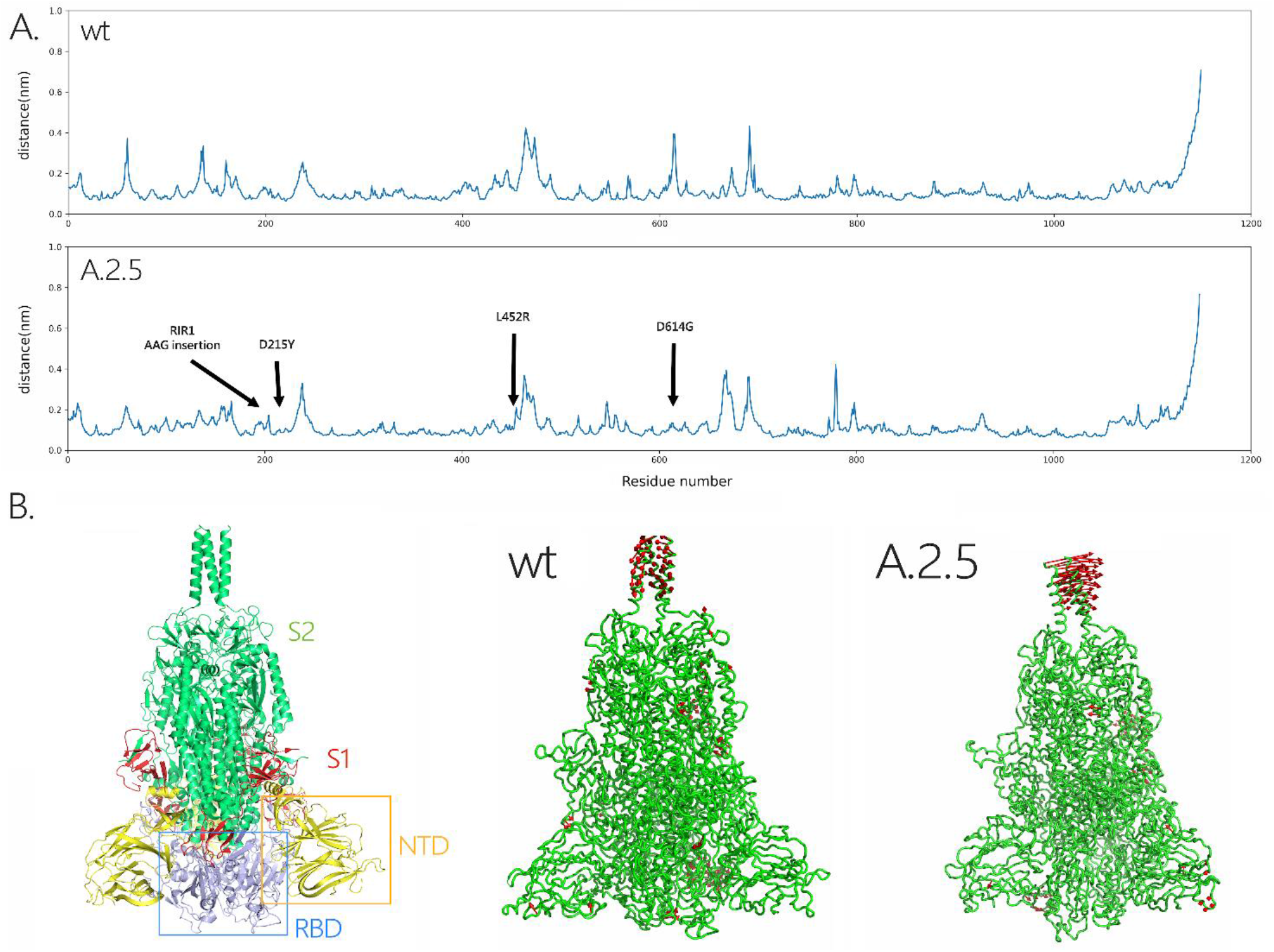
Panel A: RMSF plot for the models of the wild-type and A.2.5 SARS-CoV-2 spike proteins, with indication of the point and insertion mutations present in the two viral lineages target of his study, compared with the wild type virus. Panel B: Three-dimensional structural models obtained for the wild type and A.2.5 spike proteins. The location of the NTD and RBD (within the S1 subunit) and of the S2 subunit in the spike trimer are shown at the left-hand side. The regions where the most significant fluctuations are marked in red.

## 4. Conclusions

The SARS-CoV-2 genome continues to accumulate mutations at a relatively constant rate, occasionally originating new VOCs and VOIs as a result of continued high viral circulation and natural selection. Prior to November 2021, the insertions at RIR1 documented in this work had only led to the emergence of two viral lineages with widespread transient distribution, i.e. B.1.214.2 and A.2.5. However, the presence of a RIR1 insertion in the emerging omicron VOC, together with the recurrent independent occurrence of this phenomenon by convergent evolution in multiple viral lineages, suggests that RIR1 insertions may be linked with an evolutionary advantage, whose magnitude is presently unclear.

In absence of functional data, the role of RIR1 insertions can be only speculated. Based on the lack of overlap with known immune epitopes, their involvement in immune escape phenomena appears unlikely, even though their impact on T-cell response remains to be investigated. Similarly, the previously hypothesized role of NTD insertions in enhancing viral infectivity by promoting the interaction with host cell membranes is only partly supported by the over-representation of lysine and arginine residues in RIR1 inserts. On the other hand, we observe a correlation between the presence of RIR1 insertions, RDR deletions and several non-synonymous RBD mutations with known impact on immune evasion on infectivity. This, together with the predicted effect of RIR1 insertions on the structure of the spike protein, may suggest a possible role as a permissive mutation in compensating otherwise slightly disadvantageous non-synonymous spike mutations. Undoubtedly, our observations strongly suggest that the functional and structural impact of these insertions should be the subject of in-depth studies in the near future.

## Abbreviations

ACE2: angiotensin converting enzyme 2
NTD: N-terminal domain
RBD: receptor binding domain
RBM: receptor binding motif
RGYR: radius of gyration
RIR1: recurrent insertion region 1
RMSD: root mean square deviation of backbone beads
RMSF: root mean square fluctuations
RDR: recurrent deletion region
VOC: variant of concern
VOI: variant of interest

## Supplementary Figures

**Supplementary Figure 1.**
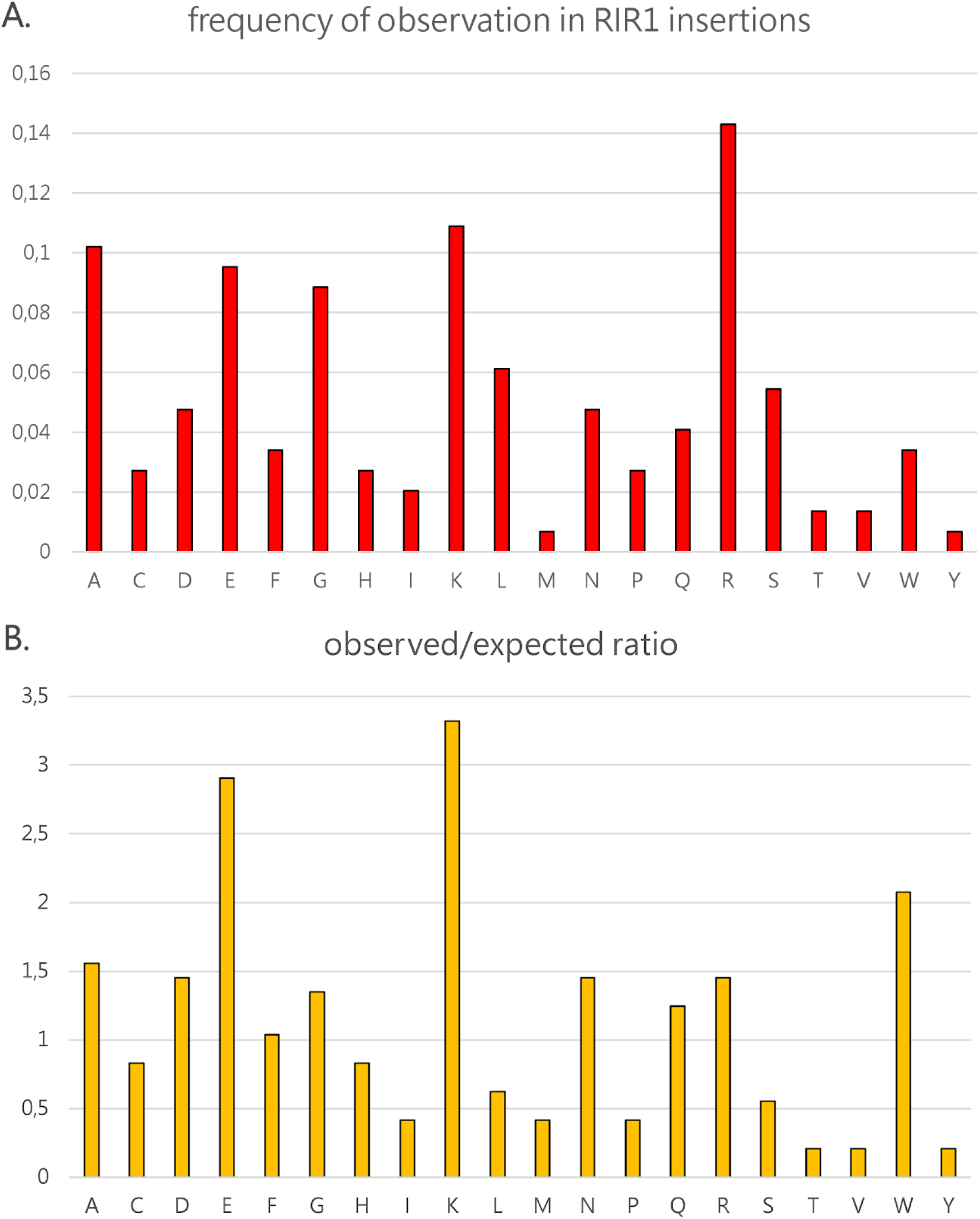
Panel A: observed frequency of each amino acid in the 41 RIR1 insertions. Panel B: observed/expected ratios for each amino acid, calculated based under the assumption that no codon usage bias was present. Amino acids showing a ratio > 1 were over-represented compared with expectations, whereas those showing a ratio < 1 were under-represented.

**Supplementary Figure 2.**
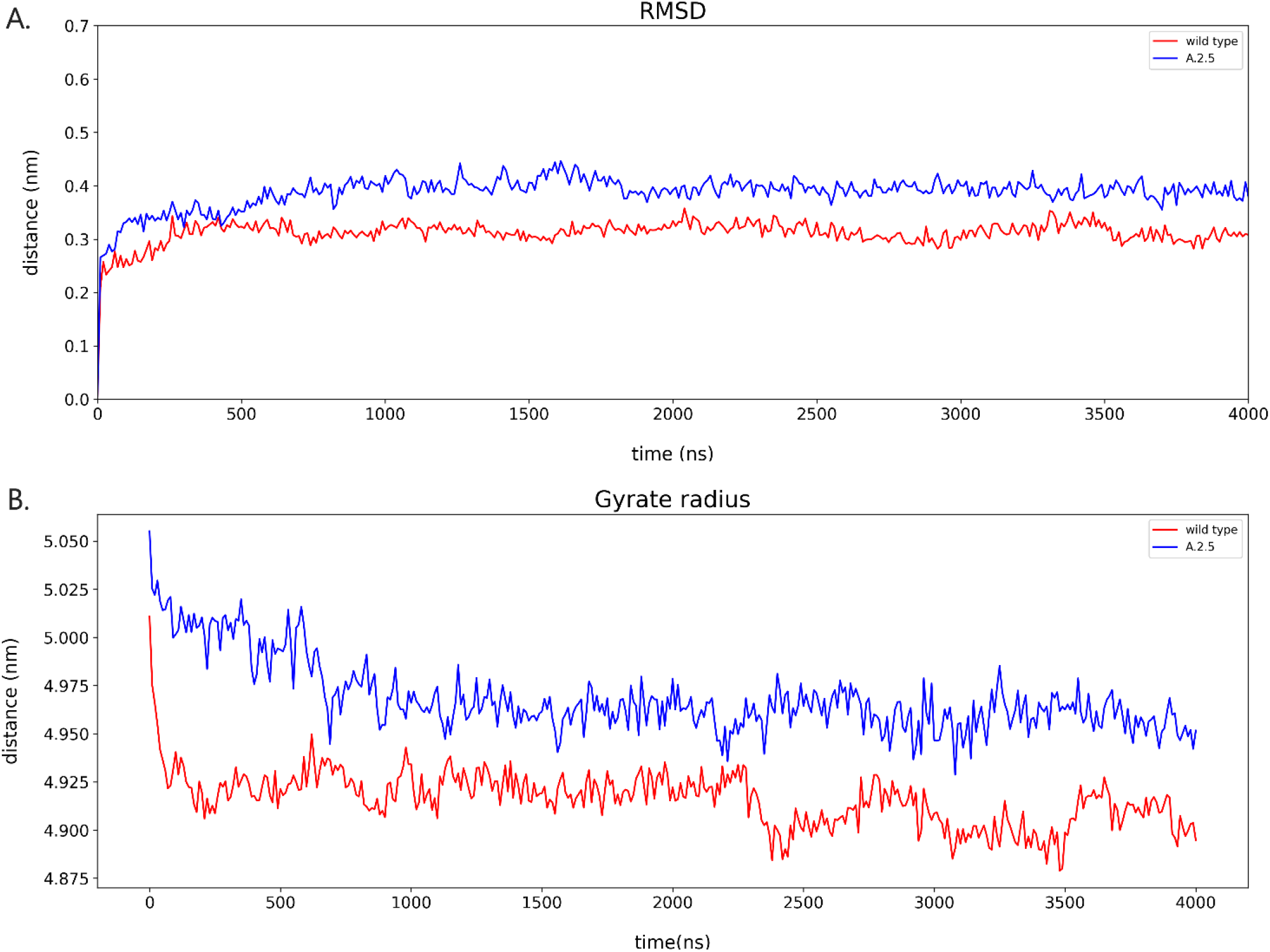
Panel A: RMSD plot of the three models of the SARS-CoV-2 spike protein (wild-type and A.2.5), as a function of simulated time. The systems reach equilibrium after 1.5 μs. Panel B: variation of RGYR observed over time for the three models of the SARS-CoV-2 spike protein (wild-type and A.2.5) during the MD simulation.

## Acknowledgements

The author is grateful to Dr. Alberto Beretta, Dr. Sarah Ann Nadeau and Dr. Alexander Martinez for their communication and useful suggestions, and the staff of Pop Medicine (https://www.facebook.com/medicipop/) for the support. The author also acknowledges all the fundamental work carried out by the clinicians, researchers and public health authorities that allowed the collection of SARS-CoV-2 genome data and made sequence data available in a timely manner though GISAID (Lopez Bernal et al., 2021), as well as the great efforts made by the developers of nextstrain to assist researchers in SARS-CoV-2 evolution studies.

## Author contributions

MG: Conceptualization; Data curation; Formal analysis; Investigation; Methodology; Supervision; Visualization; Writing - original draft; Writing - review & editing. KD, AG: Formal analysis; Investigation; Writing - original draft; Writing - review & editing.

